# Vaping Perpetuates Cardiac Electrical Instability

**DOI:** 10.1101/862441

**Authors:** Obada Abouassali, Mengmeng Chang, Michelle Reiser, Manasa Kanithi, Ravi Soni, Bojjibabu Chidipi, Sami F. Noujaim

**Author notes:** Correspondence: Dr. Sami Noujaim, Molecular Physiology and Pharmacology, Morsani College of Medicine, University of South Florida, Tampa, USA.

## Abstract

**Background:** Tobacco cigarette smoking is on the decline, but the usage of electronic nicotine delivery systems (ENDS) is gaining popularity, specifically in the teen and young adult age groups. While the cardiac toxicity of tobacco cigarette smoking has been widely studied and is well established, the possible cardiac toxicity of ENDS products and their design characteristics, such as added flavorings, are largely underexplored. Vaping, a form of electronic nicotine delivery, uses “e-liquid” to generate “e-vapor”, a smoke-like aerosolized mixture of nicotine and flavors. Here, we tested the hypothesis that vaping results in cardiac electrophysiological instability and arrhythmogenesis. We thus investigated how e-liquids with different flavors affect cardiac in-vitro and in-vivo toxicity, in cell culture and in animals.

**Methods:** Three e-liquids with vanilla, cinnamon or fruit flavors were studied. We quantified apoptosis and oxygen consumption rate in HL-1 cells cultured with e-vapors extracts. In human iPSC derived ventricular cardiomyocytes (hiPSC-CM) cultured with e-vapor extract, beating frequency and repolarization duration were measured using multiple electrode arrays (MEA). Mass spectrometry (GC-MS) was used to analyze the composition of the e-vapors. Telemetric ECGs were obtained in freely moving C57BL/6J mice exposed to vanilla flavored e-vapor for 10 weeks and heart rate variability was analyzed (HRV). In-vivo inducibility of ventricular tachycardia as well as optical mapping of voltage in isolated Langendorff-perfused hearts were also carried out.

**Results:** E-vapor caused a dose dependent increase in toxicity in Hl-1 myocytes and e-vapors containing vanilla and cinnamon flavorings, as indicated by GC-MS, were more toxic, and inhibited cellular respiration more than the fruit flavored e-vapor. In hiPSC-CM cultured with 25% cinnamon flavored e-vapor for 24 hours, beating frequency increased, and the field potential duration significantly increased compared to air control. Inhalation exposure to vanilla flavored e-vapor for 10 weeks caused significant effects on HRV. Additionally, inducible VT was significantly longer, and in optical mapping, the magnitude of ventricular action potential duration alternans was significantly larger in the exposed mice compared to control

**Conclusion:** The widely popular flavored ENDS are not harm free, and they show potential toxicity to the heart, in-vitro, and in vivo. Further studies are needed to further assess their cardiac safety profile, and long-term health effects.

## INTRODUCTION

The use of electronic nicotine delivery systems (ENDS) has been proliferating. It was recently shown that among high school students, ENDS use increased from 1.5% in 2011 to 20.8% in 2018^1^. Alarmingly, from 2017 to 2018 alone, there was a 75% increase in ENDS use by high school students^1^. The popularity of flavored ENDS likely fueled the proliferation of manufacturers, and the surge in sales of these products^2^. In 2014, it was estimated that there were more than 7600 different flavored ENDS products from 466 brands^3^, and as of this date, these numbers have only increased. The demand for ENDS continues to grow^4^ as evident by a dynamic market^3, 5^ which is projected to surpass $6 billion in the next couple of years. However, the health effects and particularly, the cardiac toxicity of ENDS remain incompletely understood.

It has been argued that ENDS use might be less harmful in some ways compared to combustible tobacco cigarettes, however, ENDS products are not harm free. Data suggests that in young people, ENDS consumption instead of the traditional tobacco cigarettes paradoxically puts the user at a significantly greater risk of later initiation of combustible tobacco cigarette smoking^1, 6^.

Vaping is a form of electronic nicotine delivery, where the vaping device heats the “e-liquid” via a coil, in order to generate “e-vapor”, an inhalable smoke-like aerosolized mixture containing nicotine and flavors. E-liquids are usually a mixture of propylene glycol and vegetable glycerin, flavors, and either nicotine salt or free base nicotine. E-liquids can be used with different ENDS devices such as the pod-based system that requires the use of e-liquid with nicotine salt, or the tank-based vaping system, where a “tank” holds the e-liquid with free base nicotine. Both pod-based and tank-based systems are popular among different age groups^4^.

Several studies investigated the toxicity of e-liquids, and it was shown that flavoring aldehydes could be harmful in cell culture^7-18^, however, the possible cardiac electrophysiological toxicity of vaping has not been systematically examined and is not completely understood. Here, we will assess the cardiac electrophysiological toxicity of 3 e-liquids of different flavors and we will test the hypothesis that vaping results in cardiac electrophysiological instability and arrhythmogenesis.

## METHODS

### Vaping chamber

A rat housing cage (Techniplast GR900) was modified, were the bottom was fitted for the introduction of the mouthpiece of Smok Species Baby V2, SMOKTech, vaping device. We used the Baby V2 A2 dual sub-coils with a total resistance of 0.2 ohms, at 85 Watts. An inlet and outlet openings were created in the cage’s lid. The inlet opening was used to connect the mouthpiece via a plastic tube (Fisherbrand ¼ ID, 3/8 OD), to an air flow meter, 4 L/ minute (Western Medica), which in turn, was connected to a silent fish tank air pump. The outlet opening was fit with a plastic tube (Fisherbrand) that served as exhaust. A Universal High-powered Door Actuator (ZoneTech)- car door locking mechanism- was fixed alongside the vaping device, aimed at the device’s firing button. The actuator was connected to an AC/DC power adapter. Both, the actuator’s power adapter and the air pump were wired into the same cycle timer. Every 2 minutes, the cycle timer turns on for 5 seconds. This causes the actuator to push the vaping device’s firing button, and simultaneously, air flows into the mouthpiece, expelling, for 4.7 seconds, an e-vapor puff at 4L/minute inside the cage. The vaping device touch screen displays the duration of the device’s activation every time the firing button is pressed. A total of 60 puffs, over a 2-hour period was delivered. This is consistent with the topography of vaping in ENDS users, where studies indicated that users could engage in an average of 50 puffs per day^19-21^. Figure 1 A is the tank-based vaping device used, and B is a diagram the exposure system.

**Figure 1:**
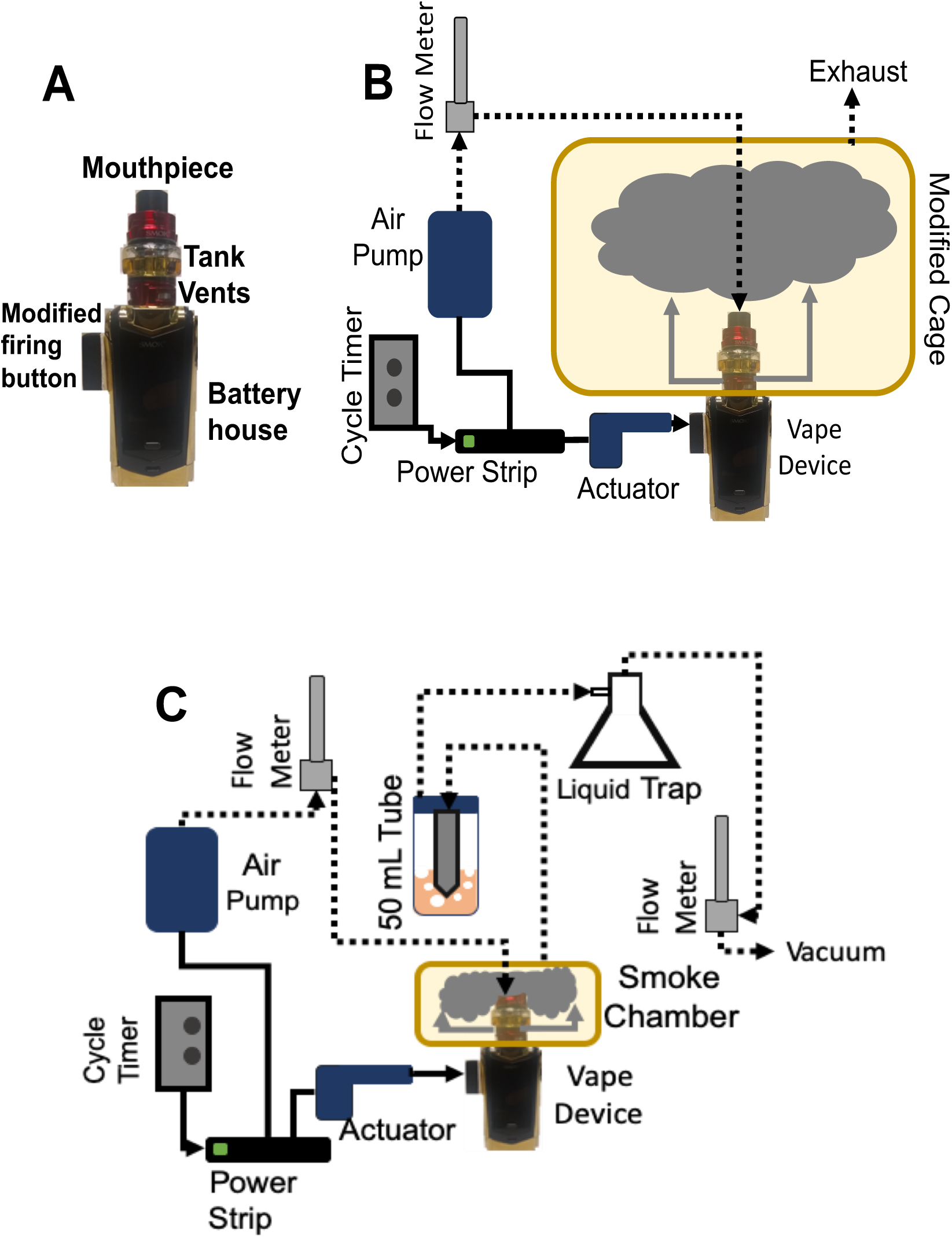
A: Picture of the Smok Species Baby V2 vaping device. The black screen displays the duration of the activation every time the firing button is pressed. B: Vaping chamber design. C: E-vapor extract generation.

### Exposure of animals to e-vapor

The animals were exposed to 4.7 second puffs of e-vapor at 4L/minute, every 2 minutes for a total of 60 puffs in a 2-hour period. Mice were exposed 5 days a week, for a period of 10 weeks. Control mice experienced the same handling of the experimental animals and were placed in a similar modified chamber in the same environment where the only difference is that they were exposed to normal room air.

### Preparation of e-vapor extracts

A 10cm × 10cm × 7cm chamber was modified with a bottom opening fitted for the mouthpiece of the Smok Species Baby V2, SMOKTech, vaping device, and an inlet and outlet openings were introduced in the chamber’s sealing top lid. The inlet tube was connected by plastic tubing (Fisherbrand) into the vaping device’s mouthpiece and the flow meter and air pump, as done for the vaping cage above. The outlet was passed through the cap of a 50 ml conical tube containing cell culture media. Another tube was passed into the 50 mL conical tube cap and connected to a liquid trap flask, then to an air flow meter at 4L/ minute, and finally to a vacuum. Similar to the vaping machine, every 2 minutes, the cycle timer turns on for 5 seconds causing the actuator to push the vaping device’s firing button, and simultaneously, air flows into the mouthpiece, expelling a 4.7 seconds e-vapor puff at 4L/minute inside the chamber. The vacuum pump draws the e-vapor from the chamber at 4L/minutes, leading to its bubbling into the medium. 10mL of medium was bubbled with 15 puffs of vapor, considered to be 100% e-vapor extract. For air control, the exact procedure was performed, however, the vaping device was powered off. Dilutions of e-vapor, or air extracts were then made with non-treated cell culture media. Figure 1C is a diagram of the e-vapor extract system.

### E-liquids

The flavors used in this study were, Hawaiian POG (POG), and Vanilla Custard (VC) by USA Vape Labs, CA, and Apple Jax (AJ), by Epic Juice, LLC. The manufacturer labeled POG flavor as passion fruit, orange and guava, Vanilla Custard, as vanilla custard, and Apple Jax as milky cinnamon apple cereal. These e-liquids are 70% vegetable glycerin/ 30% propylene glycol (70VG/30PG) and stated to contain 6mg/ml free base nicotine.

### HL1 cell culture

HL-1 cells were obtained from the laboratory of Dr. Claycomb (Louisiana State University), and cultured following the recommended protocol^22^. Briefly, cells were grown in Claycomb medium (Sigma) supplemented with 10% FBS (Sigma), 0.1 mM norepinephrine (Sigma), 2mM L-Glutamine (Sigma), and penicillin/streptomycin (100U/ml/100µ/mL) on tissue culture plates (Corning), coated with fibronectin/gelatin (Sigma).

### Apoptosis flow cytometry

Apoptosis was measured using the FITC annexin V staining assay (BD Pharmingen). HL-1 cells were plated in 12 well plates, and when confluency reached approximately 70%, they were cultured with air bubbled control medium, or with medium with e-vapor extracts. After the incubation period, the cells in each well were lifted with Accutase and re-suspended in 5 mL PBS. Cells were pelleted by centrifugation, and washed with PBS and stained with FITC labeled annexin V according the manufacturer’s recommendation. DAPI (3 µM) was added immediately before reading the samples on a BD LSRII Cytometer using the 488-nm and 405-nm lasers for excitation of FITC annexin V and DAPI respectively. The following controls are included in each experiment: unstained cells, cells stained with FITC annexin V, but not with DAPI, and cells stained with DAPI, but not with FITC annexin V. Data analysis was carried out using the FlowJo software.

### Human induced pluripotent stem cells derived cardiomyocytes culture and extracellular potentials recording

The iCell Cardiomyocytes2 (Fujifilm) were thawed according to the manufacturer’s protocol, and 50,000 cells/well are plated on the fibronectin (50 µg/ml) coated 24 wells multiple electrode array (MEA) plates (24well plate-eco, MED64). Plated cells were incubated at 37°C, 5% CO2 for one hour to allow attachment of the cells, after which, pre-warmed maintenance medium (Fujifilm) was added to the wells. The maintenance medium was changed every three days, and after day 7, cells began to beat synchronously, periodically and spontaneously. The Presto Multielectrode Array (MED64) was used to record extracellular potentials of the spontaneously hiPSC derived cardiomyocytes. The Presto is equipped with an environmental chamber (37°C and 95% O2/ 5% CO2). After the plate was placed on top of the recording electrodes, 2-minute recordings of the extracellular potentials were obtained using the MED64’s MEA Symphony software interface at base line before addition of the vapor extracts, then 24 h and 48 h after addition of the extracts. Data was filtered offline with a 200 Hz low pass Butterworth filter and a 0.1 Hz high pass Butterworth filter. For each well, the beating rate, and the field potential durations were calculated in the MED64’s MEA Symphony analysis software.

### Mass spectrometry

E-vapors were generated using the same setup described in Figure 1. The 10cm × 10cm × 7cm chamber was slightly altered, where the outlet tube was replaced with a septum. 5 puffs were generated, each for 4.7s at an interval of 10s, at a flow rate of 4 L/min. Immediately after the smoke was generated, a 2.5mL Hamilton glass syringe was introduced through the septum to extract 250uL of e-vapor from the chamber. The smoke was then immediately injected manually into the Aglient 7890B GC-MS 5977B. The headspace syringe was cleaned between sample by rinsing repeatedly in pesticide grade methanol and then thoroughly dried in an incubator for 30mins at 70 C. The GC-MS parameters were optimized based on previous studies^17, 23^ and are reported in Table 1. MassHunter Workstation Qualitative Analysis Software (Version B.07.00 SP2) in conjunction with NIST MS Search 2017 Library wereu sed for analysis. Peaks were identified based on a match score factor higher than 700.

**Table 1:** GC-MS settings used for qualitative analysis of the e-vapors.

| **GC-MS Parameter** |  |
| --- | --- |
| Capillary Column (GC) | HP-5MS UI (30m x 250 mm x 0.25μm) |
| Helium Flowrate (GC) | 1.00 mL/min |
| Inlet Temp. (GC) | 250°C |
| Oven Program (GC) | Ramp 1: 40°C to 170°C @ 10°C/min; hold 2 min; Ramp 2: 8°C/min to 250°C; Ramp 3: 25°C/min to 320°C; hold 5 min. |
| Split Ratio (GC) | 20:1 |
| Auxiliary Temp. (MS) | 290°C |
| Solvent Delay (MS) | 1.00 min |
| Total Ion Scan (MS) | m/z = (30-450) from 1 min to 32.80 min |

### Oxygen consumption rate measurement

The oxygen consumption rate (OCR) of HL-1 cardiomyocytes was measured using the XFp Cell Mito Stress Test by Agilent Seahorse^24^. Modulators of cellular respiration are employed to quantify ATP-linked respiration with 1.5 μM of oligomycin, maximal respiration, with carbonyl cyanide-4-(trifluoromethoxy) 1.5 μM phenylhydrazone (FCCP), and nonmitochondrial respiration driven by processes outside the mitochondria with 0.5 μM Rotenone, and Antimycin. Two days prior to the experiment, 40,000 cells/well were plated in a gelatin/fibronectin precoated XFp mini cell culture plate (11.4 mm2 area) at 37 □°C. Cells were treated with 25% e-vapor extract for 24 h. Right before the measurement, the culture medium was replaced with the prewarmed assay medium (Agilent) containing (in mM) Glucose 10, Pyruvate 1, and L-Glutamate 200. Oligomycin, FCCP, and Rotenone/Antimycin are added to the cells in 20 minutes intervals, according to the essay’s manufacturer’s instruction in order to measure ATP linked, maximal, and non-mitochondrial oxygen consumption rates^24^. Soon after the experiment was completed, media was gently removed, and 40µl of RIPA lysis buffer was added, and cells were lysed using a 1 ml syringe. Protein concentration was quantified by BCA protein assay (Thermo Fisher Scientific) for normalization of the respiration parameters.

### Telemetry and HRV analysis

Wild type C56bl/6J (males and females,3 to 5 months of age), were implanted with ECG telemetric devices (ETA-F10, DSI) using sterile equipment, under general anesthesia with 2% Isoflurane, and body temperature maintained at 37°C on with a heating pad. Manufacturer’s recommendation for subcutaneous transmitter placement along the lateral flank were followed. The animals were allowed to recover for fourteen days after the implantation procedure. ECG recordings in freely moving animals, were done using the PhysioTel RPC-3 (DSI) receivers, after 1, 5, or 10 weeks of exposure to e-vapor. Two minutes ECG strips were analyzed in the Lab Chart 8 (AD Instruments) HRV analysis software as previously done for mice^25^. In the Fast Fourier Transforms, very low frequency was defined as 0hz to 0.15hz, low frequency as 0.15 Hz to 1.5 Hz, and high frequency as 1.5 Hz to 5 Hz similarly to what is done in mouse HRV studies^25, 26^

### In vivo VT inducibility

Mice were anesthetized (2% isofluorane), and a 1.2 French octapolar catheter (Millar) placed transvenously into the right atrium. Electrograms were recorded using the PowerLab platform from Advanced Instruments. Programmed electrical stimulation for VT induction was performed at twice diastolic threshold with 1 second burst pacing from 20 to 50 Hz, in 2 Hz increments^25, 27^.

### Optical imaging

Isolated Langendorf perfused mouse hearts were retrogradely perfused with Tyrode’s solution. The preparations were maintained at 37°C and stained with a bolus of voltage sensitive dye (0.25 ml, 10 μM Di 4 ANEPPS, Molecular Probes). Blebbistatin (7 μM, Tocris) was used as an excitation-contraction uncoupler. Mapping was carried out as we have done extensively. We quantified the action potential duration at 75% repolarization (APD) as we previously did^28, 29^. A bipolar, silver tip stimulation electrode was used to pace the ventricles (5ms pulses, 2x diastolic threshold) at different frequencies (from 8 to 15 Hz).

### Statistics

Data are presented as average ± standard error. Student’s t-test, one way analysis of variance (ANOVA) with Bonferroni correction, or Kruskal-Wallis test with multiple comparisons were used as appropriate and significance was taken at p<0.05.

## RESULTS

HL-1 mouse atrial cardiomyocytes^22^ were cultured for 48 h with 5, 10, 25 and 50% air or with vanilla custard (USA Vape Lab, CA) e-vapor extract. Cells were then dissociated with Accutase and stained with FITC labeled annexin V, a widely used apoptotic marker, and DAPI, a cell viability marker. Figure 2 A, left panel, shows a flow cytometry analysis of air controls. The majority of cells are in the low annexin V and low DAPI live quadrant. Apoptotic cells show high annexin V (right lower and upper quadrants). Necrotic cells show low annexin V and high DAPI labeling. In vanilla custard e-vapor treatment, there was a significant shift of the population, from the live quadrant, to the necrotic and apoptotic quadrants. Panel B is a quantification of the percentage of live cells at 5, 10, 25 and 50% treatment with air, or vanilla custard e-vapor. There was a dose dependent decrease in the live population as the concentration of vanilla custard e-vapor increased. The percentage of necrotic (panel C) and apoptotic (panel D) cells increased with increasing vanilla custard e-vapor extract. Panel E is a plot of the toxicity index (TI) which we calculated as 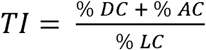 where %DC is percentage of dead cells, %AC is the percentage of apoptotic cells, and %LC is percentage of live cells per well. The TI was normalized to that of air at each treatment condition. Figure 2E is the TI normalized to that of air at 5, 10, 25 and 50% vanilla custard extract, where TI was 1.3 ± 0.04, 1.4 ± 0.03, 4.3 ± 0.13, and 13.6 ± 3.5 respectively.

**Figure 2:**
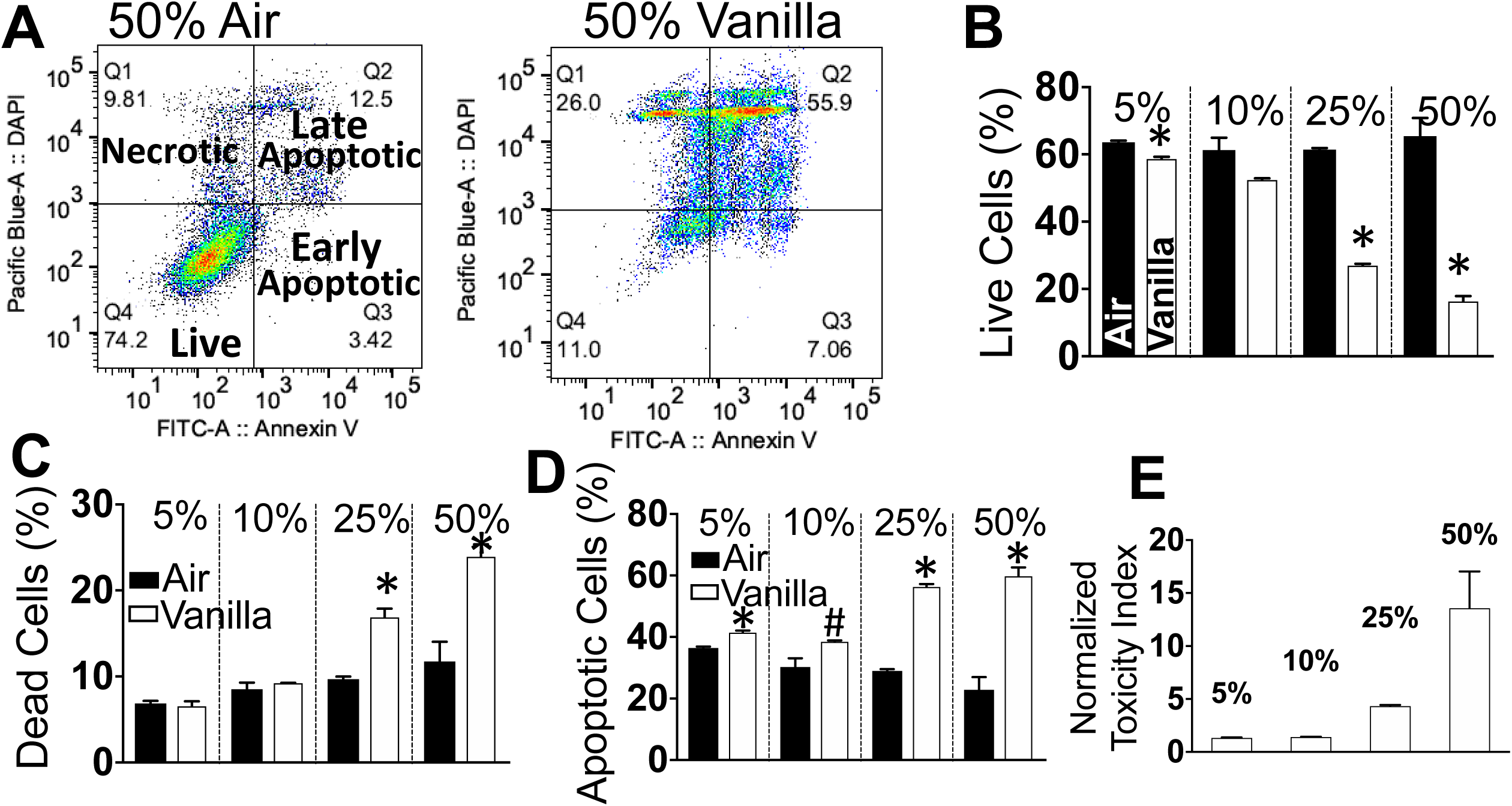
Quantification of Annexin V staining in HL-1 cells with flow cytometry. A: Flow cytometry analysis of live, apoptotic, and necrotic HL-1 cells cultured for 48 h with 50% air (left), or 50 % Vanilla Custard e-vapor extract. B,C,D: Flow cytometry quantification of the percentage of live (B), necrotic (C), and apoptotic (D) cells, in air or Vanilla Custard e-vapor bubbled medium at 5, 10, 25 and 50%. E. Toxicity index normalized to that of air. N= 3 in each condition.

Figure 3A compares the effects of 48 h, 50% e-vapor extracts of Hawaiian POG (POG), vanilla custard (VC) and apple jax (APJ) or air on HL-1 myocytes. Flow cytometry of annexin V and DAPI staining showed that all three e-liquids decreased viability (top panel) and increased apoptosis (middle panel) to different extents (*p<0.01, vs air). The bottom panel shows that vanilla custard and apple jax were more toxic compared to POG (*p<0.01). We then used gas chromatography mass spectrometry (GC/MS) to analyze the chemical composition of these e-liquids. Figure 3 B shows the chromatograms of vanilla custard (VC), apple jax (APJ) and POG. The complete list of identified constituents in these e-liquids is presented in Table 2. The propylene glycol peak was trimmed to enhance visibility of the smaller peaks of flavorings as done elsewhere^17,23^. The glycerin and nicotine peaks are evident. In vanilla custard and apple jax, but not in POG, flavorings peaks corresponding to aldehydes products such as cinnamaldehyde (red arrow), vanillin, and ethyl vanillin (green arrows) are present. This is consistent with reports in cell culture, showing that vanilla, and cinnamon containing flavors cause higher toxicities^17^.

**Table 2:** GC-MS peaks analysis of the e-vapors.

| Vanilla Custard | Apple Jax | Hawaiian POG |
| --- | --- | --- |
| 2(3H)-Furanone, dihydro-5-pentyl- | 2-Propenoic acid | 2(3H)-Furanone, 5-hexyldihydro |
| Acetoin | 2(3H)-Furanone, 5-hexyldihydro- | 3-Hexenol |
| Anisyl alcohol | Benzyl alcohol | Cinnamic acid, methyl ester |
| Benzaldehyde, 4-methoxy | Butanoic acid, 3-methyl-, ethyl ester | D-Limonene |
| Benzene, 1,4-dimethoxy | Cinnamaldehyde | Furaneol |
| Butanoic acid, 3-methyl-, 3-methylbutyl ester | Coumarin | Glycerin |
| Butanoic acid, ethyl ester | Cyclopentanol, 1-(1-naphthalenyl)- | Maltol |
| Cyclotene | Ethyl maltol | Proplene glycol |
| Ethyl maltol | Ethyl Vanillin | Pyridine |
| Ethyl Vanillin | Glycerin |  |
| Glycerin | Maltol |  |
| Phenol, 2-methoxy | Piperazine |  |
| Propylene Glycol | Propylene Glycol |  |
| Pyridine | Pyridine |  |
| Sulfurol | Vanillin |  |
| Vanillin | Vanillin propylene glycol acetal |  |
| Vanillin propylene glycol acetal |  |  |

**Figure 3:**
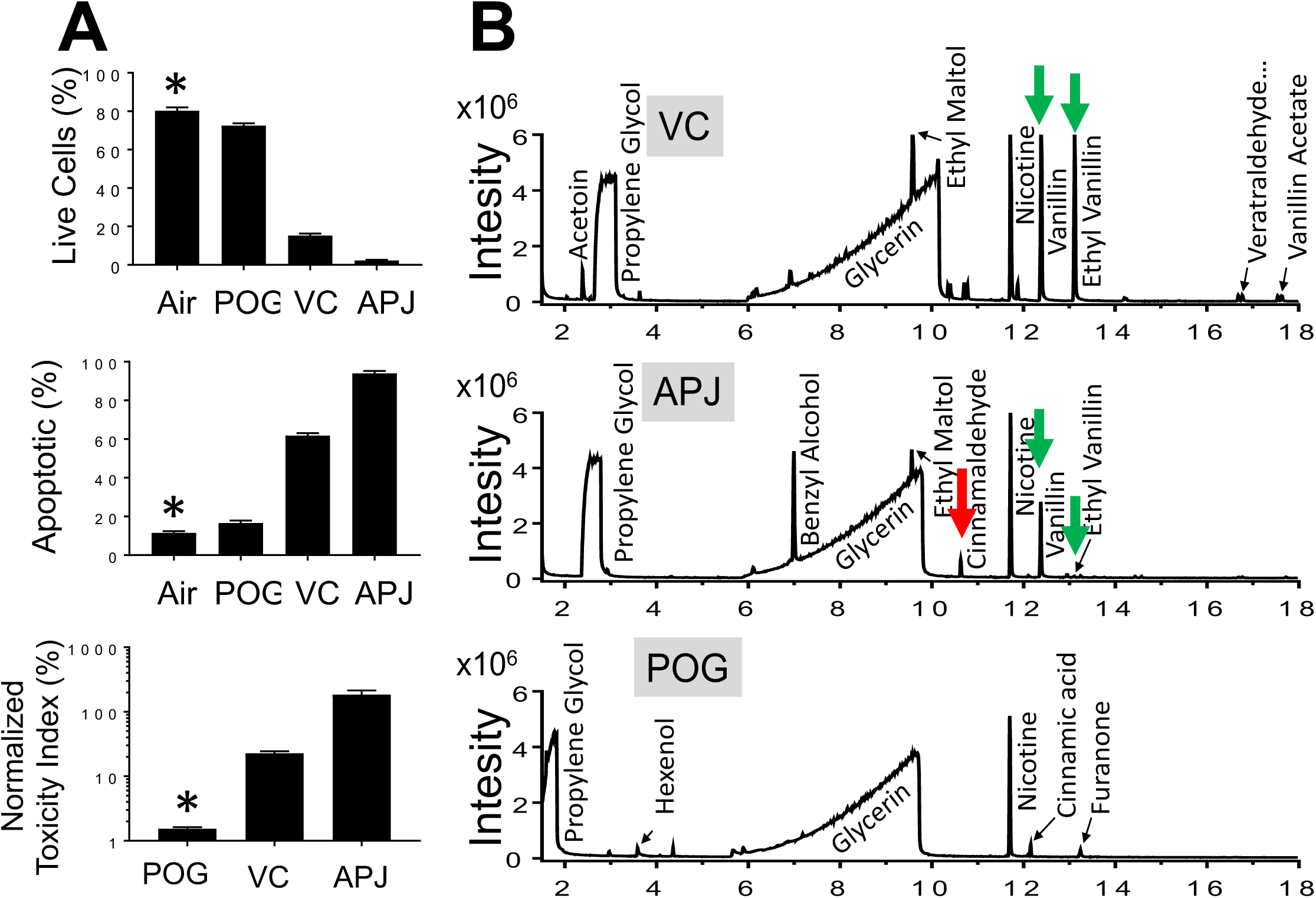
A: Toxicity of Air, POG, Vanilla Custard (VC), and Apple Jax (AJ) e-liquid vapor extracts at 50%, for 48h in HL1 cells using flow cytometry and Annexin V staining. N=3 in each condition. B: GC-MS chromatograms of vanilla custard (VC), apple jax (APJ) and POG. Propylene glycol, glycerin and nicotine peaks are evident in addition to flavorings peaks corresponding to aldehydes products such as cinnamaldehyde (red arrow), and vanillin and ethyl vanillin (green arrows). The list of identified constituents is in Table 2.

We compared cellular respiration in HL-1 cells after 24h treatment with 25% e-vapor from an e-liquid with a higher toxicity (apple jax, APJ) or lower toxicity (POG) as determined in Figure 3. Oxygen consumption rate (OCR) was measured by the Agilent Seahorse^24^. Figure 4A is a graph of OCR versus time in 25% air, APJ or POG treated cells. ATP-linked respiration was inhibited by addition of 1.5 μM oligomycin^24^ (Oligo) then maximal respiration occurred after injection of 1.5 μM phenylhydrazone^24^ (FCCP). The OCR remaining after addition of 0.5 μM rotenone + antimycin (A/R) is due to non-mitochondrial respiration^24^. Panel A curves indicated that apple jax depressed oxygen consumption more significant than air or POG (Kruskal-Wallis test with multiple comparisons, p<0.01). Panel B quantifies basal OCR (OCR before addition of Oligo), ATP linked OCR (OCR after FCCP minus OCR after Oligo), and maximal OCR (OCR before A/R minus after A/R). Basal and ATP linked OCR were significantly depressed with apple jax and POG treatment versus air, but maximal OCR was significantly lower in air versus apple jax only (one way ANOVA, Bonferroni correction, p<0.05). As a whole, this experiment suggests that flavored e-liquids impair the cardiac myocyte’s respiratory process^24^ and an e-liquid with a higher toxicity depresses the respiratory processes more that an e-liquid with a lower toxicity.

**Figure 4:**
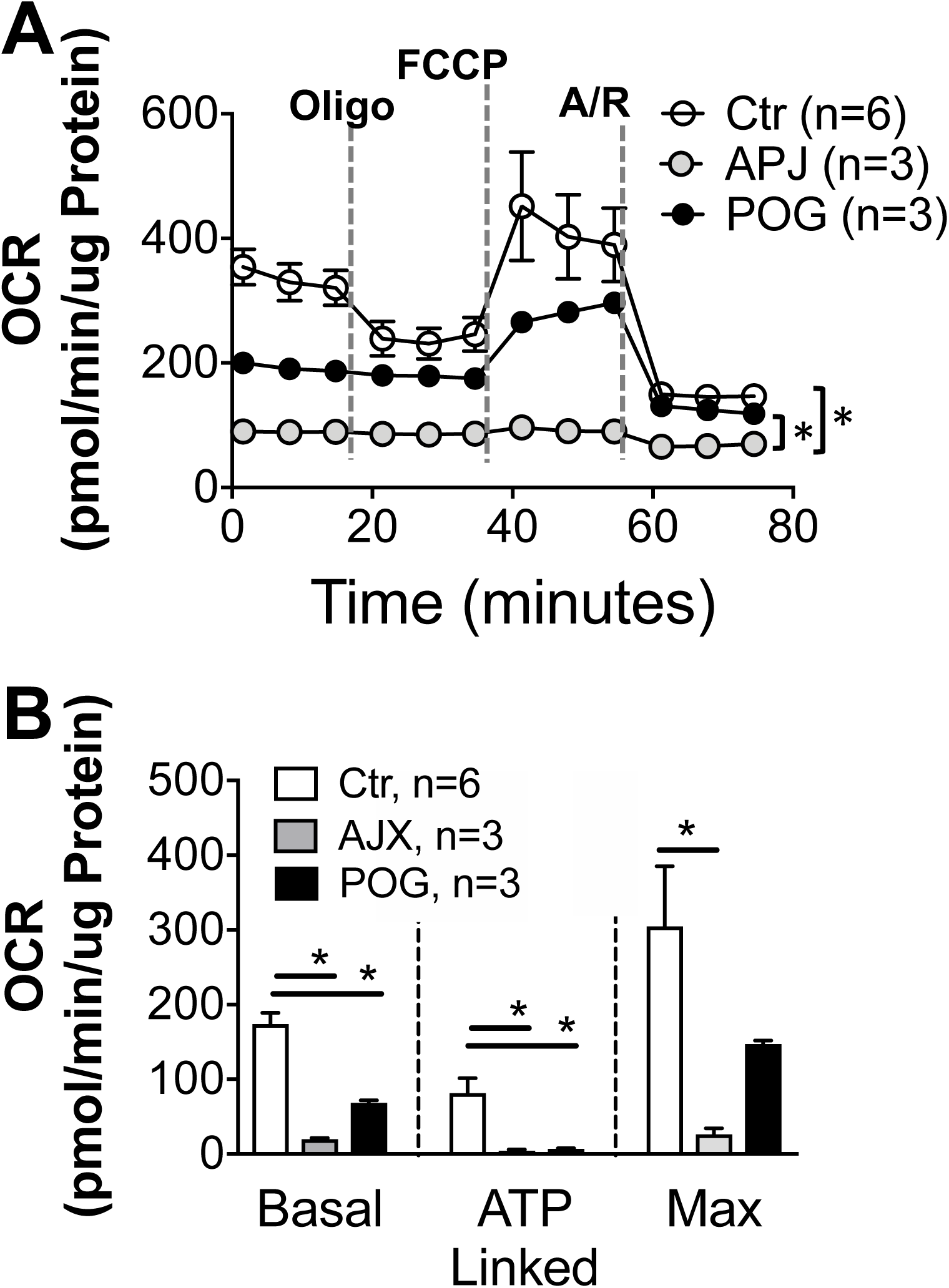
Effects of apple jax and POG e-vapors on oxygen consumption rate (OCR) in HL-cells. A: Plot of normalized OCR versus time in HL-1 cells cultured for 24 hours with 25% air or 25% apple jax or POG e-vapor extracts. The curve shows the change in OCR after injections of Oligo (oligomycin), FCCP (phenylhydrazone) and A/R (Antimycin and Rotenone). B: Quantification of normalized basal, ATP linked and maximal OCR.

Next, we tested in spontaneously beating hiPSC derived ventricular cardiomyocytes, the effects of 5, 25 and 50% apple jax e-vapor extract or air on beating rate and field potential durations. hiPSC cardiomyocytes were seeded in 24 well multiple electrode array (MEA) plates. Each well contains 16 electrodes in a 4×4 configuration, 100 μm interelectrode distance. The Presto Multielectrode Array was used to simultaneously record the 16 unipolar extracellular potentials in each well. Only wells that showed spontaneous activity were used. 50% Apple Jax e-vapor completely suppressed the spontaneous beating of the cells, but not 50% air. Figure 5 A shows the electrograms recorded before and after treatment with 25% Apple Jax e-vapor for 24 hours. 25% apple jax treatment for 24 hours accelerated the beating rate from 27 ± 1 in air control, to 60 ± 4 beats per minutes in apple jax (*p<0.01), and increased the field potential duration from 152.1 ± 10.1 ms in air to 270.6 ± 13.064 ms in e-vapor treated cells (*p<0.01). 5% apple jax e-vapor did not significantly affect beating rate or the field potential duration.

**Figure 5:**
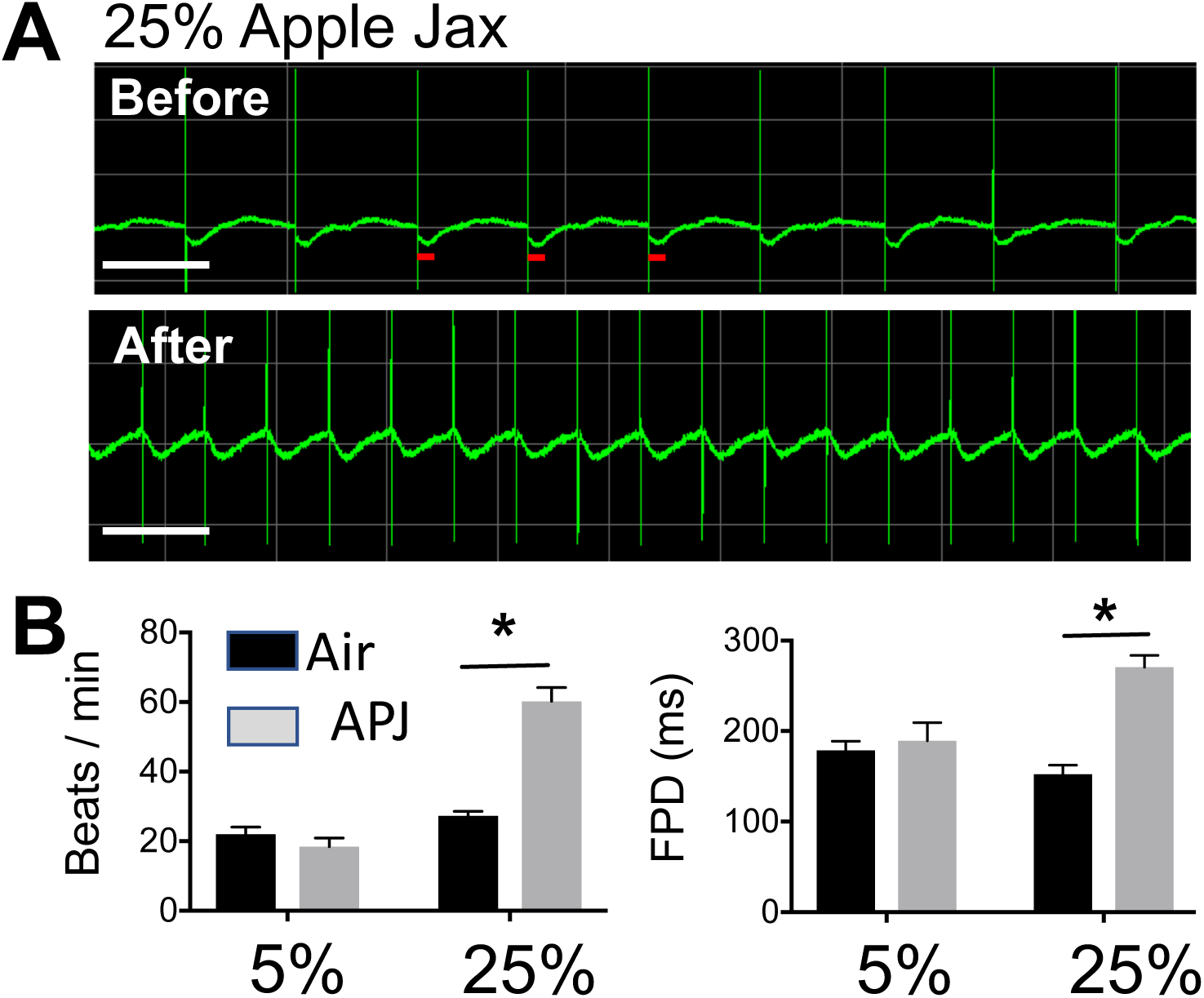
Effects of apple jax e-vapor on beating rate and field potential duration in human iPSC derived cardiomyocytes. A: Multiple electrodes array (MEA) recording of extracellular potentials in the spontaneously beating myocytes before, and after 24 hours of incubation with 25% e-vapor. B: quantification of beating rate and field potential duration (FPD) after 24 hours culture with air, or 25% e-vapor medium. N=3 wells each. White scale bar: 2 seconds.

We investigated the effects of 10 weeks inhalation exposure to vanilla custard e-vapor, or air control, on heart rate variability parameters in wild type C56bl/6J mice instrumented with telemetric ECG. HRV parameters were measured at baseline, 1, 5, and 10 weeks exposure. The RR intervals were not different in the two groups at any of the time points. In Figure 10 A, the low frequency (LF)nu component was significantly increased after 5 and 10 weeks of vaping compared to baseline (*p<0.05, One way ANOVA with Bonferroni correction). In Panel B, the high frequency (HF)nu component was significantly decreased at 5 and 10 weeks compared to baseline. In Panel C, the LF/HF ratio significantly increased at 5 and 10 weeks of vaping compared to baseline. None of the parameters changed in air control mice. The HRV changes in vaped mice are consistent with what has been shown in human subjects who use tobacco cigarettes and electronic cigarettes^30^. Such changes in HRV are indicative of sympathovagal disbalance in the control of heart rate which is clinically linked to poor cardiovascular outcomes^31^. Next, we studied the in vivo inducibility of VT in 5 wild type C56bl/6J mice exposed to vaping with vanilla custard for 10 weeks versus 7 mice exposed to air control, using in vivo programmed electrical stimulation. The ECGs of Figure 6B show that a 1 second, 36 Hz burst stimulus at 1 mA, induced a short lived ventricular tachycardia (VT) episode, after which the heart reverted back to sinus rhythm. In the vaped mouse, a similar burst stimulus induced a longer VT episode. Six out of 7 air control mice had VT, while 5 out of 5 vaped animals had VT, however, as shown in the bar graph, the duration of VT episodes was significantly longer in vaped compared to WT mice (t-test, p<0.05).

**Figure 6:**
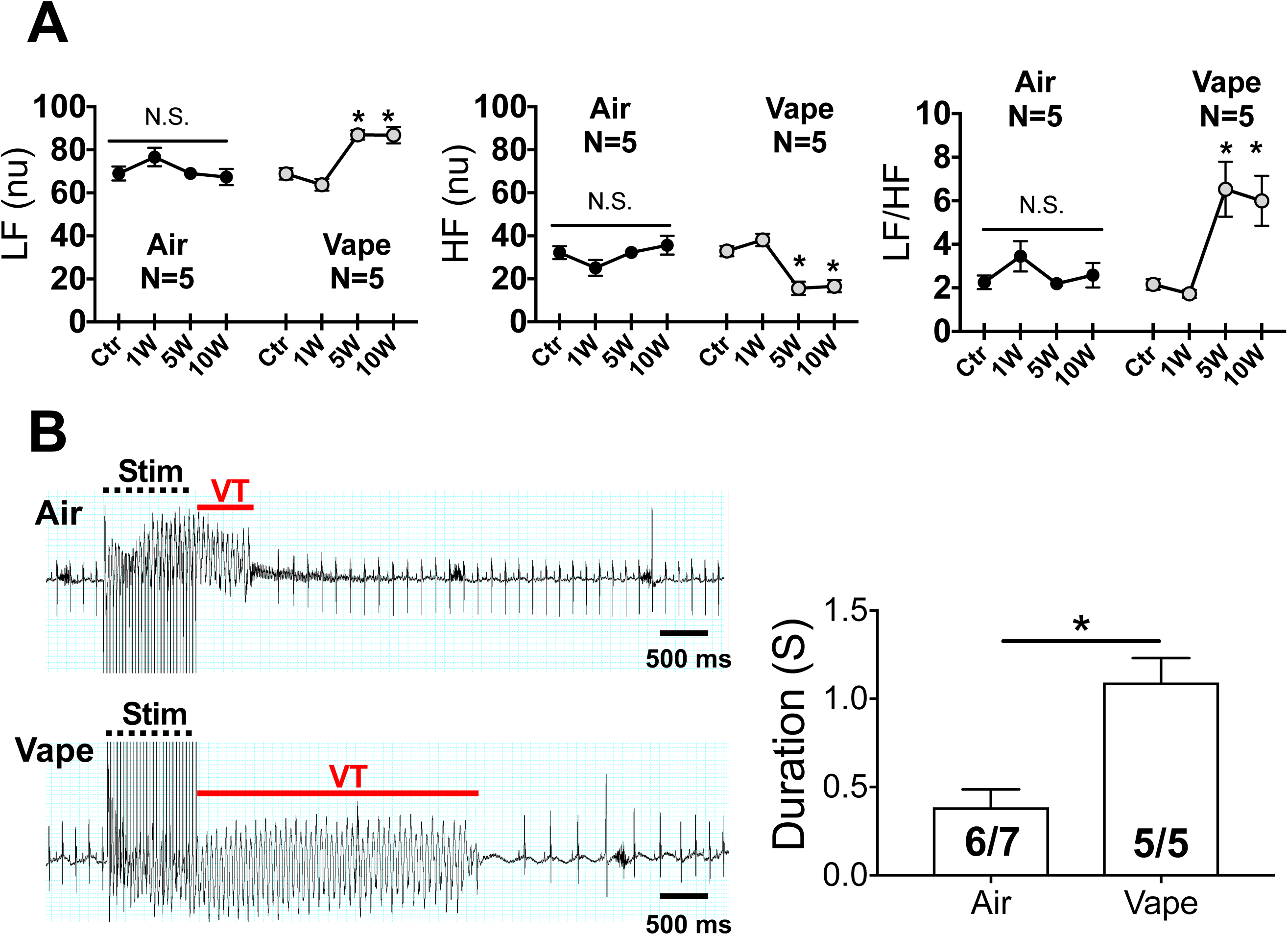
Heat rate variability (HRV) and in vivo inducibility of VT in mice exposed to air or vanilla custard e-vapor. A: Low frequency (LF)nu, high frequency (HF)nu, and ratio of low to high frequency (LF/HF) in mice exposed to air or vaping at baseline, and after 1 week (1w), 5 weeks (5W), and 10 weeks (10W). B: ECG traces of inducible VT in conrol and vaped mice. The bar graph compiles the duration of inducible VT episodes in 6 out of 7 control and 5 out of 5 vaped mice.

**Figure 7:**
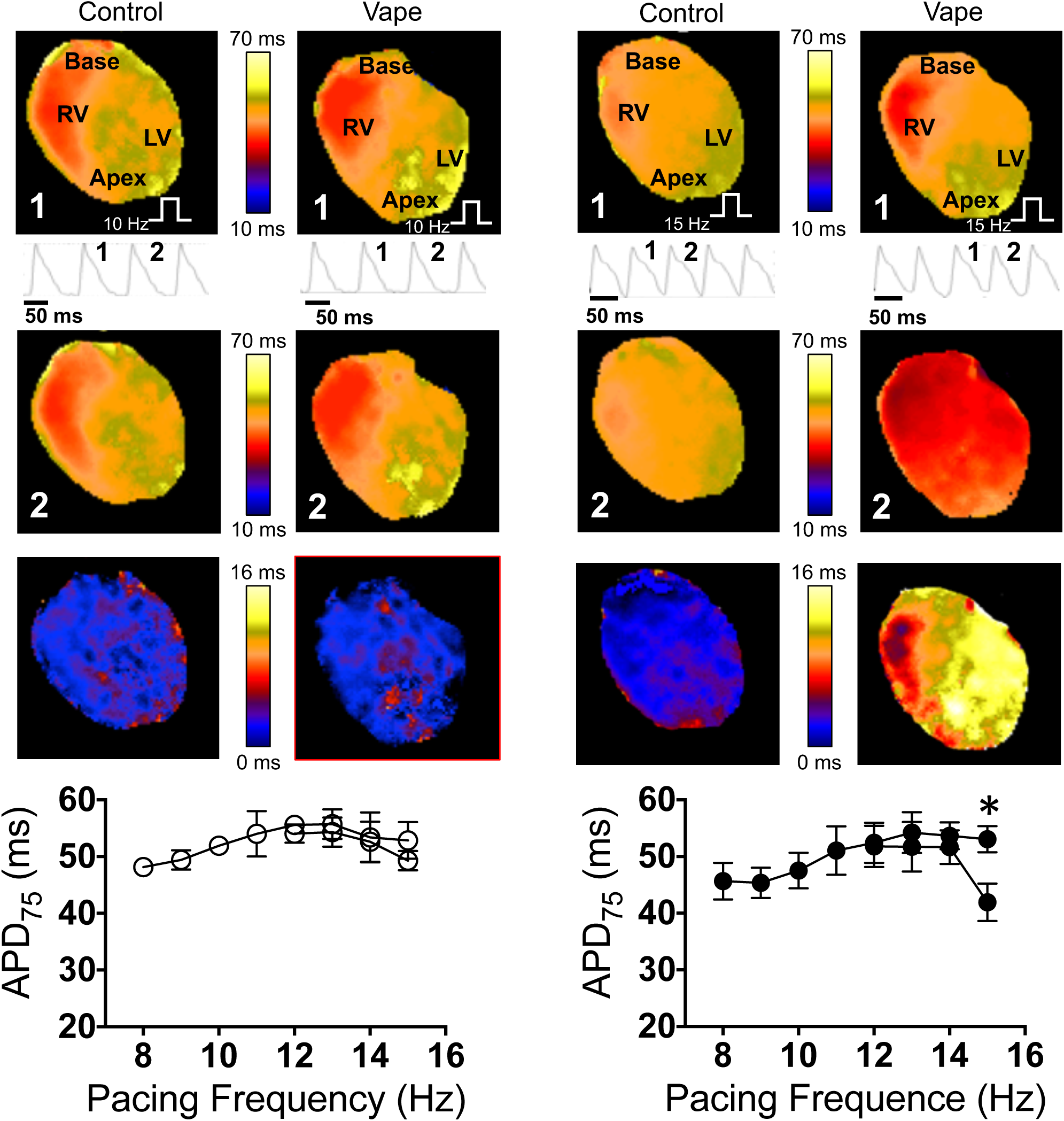
Optical mapping in isolated langendorff- perfused mouse hearts. First (beat 1) and second (beat 2) rows: APD_75_ maps of 2 consecutive beats in pacing at 10 (first 2 columns) and 15 Hz. Single pixel optical action potential traces are shown. Thrid row: difference maps of APD_75_ maps of beats 1 minus beat 2. Bottom panels: average APD_75_ at different pacing frequencies (8 to 15 Hz). RV; right ventricle. LV; left ventricle.

Subsequently, in 4 air control and 3 vaped mice we conducted epicardial optical mapping of voltage. Figure 12 panels A and B show maps of action potential duration at 75% repolarization (APD_75_) of the anterior surface of the ventricles at 10 and 15 Hz pacing respectively. Pacing was done from the apex as indicated by the white stair symbol. RV and LV are the right and left ventricles. In A, the top maps are representatives of paced beat 1, and the middle maps are representatives of the subsequent paced beat 2 in control and vaped hearts. Single pixel recordings of the action potentials are underneath the top maps, and they show beat 1 and beat 2. The action potential duration was not different between beat 1, and the subsequent beat 2, as evidenced by the difference map in bottom (map 1 minus map 2). In panel B, the hearts were paced at 15 Hz. It can be appreciated from the single pixel recordings of the optical action potential, and from the APD maps, that while in control the APD did not change considerably between beat 1 and 2, action potential duration alternans occurred in the vaped heart, where one beat is long (beat 1), while the successive beat is short (beat 2). These APD alternans can be visualized in the difference map (bottom). Panel C plots the averaged action potential durations in 4 control hearts, quantified from the APD maps at different pacing frequencies (from 10 to 15 Hz). No significant alternans occurred. Panel D is the averaged action potential durations in 3 vaped hearts, paced from 10 to 15 Hz. Significant alternans occurred at 15 Hz (*p<0,01, t-test), indicating that vaping induces action potential changes in the heart. It has been shown that alternans are indicators of cardiac electrophysiological instability and could lead to arrhythmogenesis^32^.

## DISCUSSION

Flavored ENDS are very popular, and it has been argued that the appeal of flavored ENDS products has fueled the rapid and spectacular growth of this industry^3, 5^. Thus, our objective was to investigate the in vitro and in vivo cardiac electrophysiological toxicity of flavorings in vaping. Our study showed that vaping results in cellular and whole organ electrophysiological toxicity that includes action potential instability and reduction in heart rate variability parameters. Therefore, although ENDS could be less harmful than traditional cigarette^33^, they are not harm free to the electrophysiological function of the heart.

In HL-1 mouse atrial cardiomyocytes^22^ the 3 e-liquids tested were toxic, however to different extents, where aldehyde containing vanilla custard and apple jax flavors, as determined by GC-MS, were more detrimental compared to the fruity flavored e-liquid. This is independent of nicotine content, since these e-liquids are stated to contain 6 mg/ml nicotine. These findings are consistent with similar studies conducted in many other cell lines, however, these cell lines are not relevant to cardiac electrophsyiology^11, 13-15, 17, 18^. Additionally, vaping reduced cellular respiration in HL-1 cells, where basal, ATP linked and maximal rates of oxygen consumption were significantly reduced, similarly to what has been reported with tobacco cigarette smoke in airway smooth muscle cells^34^. In spontaneously beating hiPSC derived ventricular cardiomyocytes, e-vapor increased the beating rate as well as the corrected field potential duration (a surrogate for the QTc interval)^35^, raising the possibility that vaping could be arrhythmogenic since in the heart, prolongation of the QT interval is associated with the development of fast ventricular rhythms that include tachycardia and fibrillation^36^.

HRV analysis revealed patterns of change indicative of sympathovagal disbalance in the control of the mouse heart due to vaping where LF was increased, HF was decreased and LF/HF ratio increased. These changes are in line with what has been shown in human subjects who use tobacco cigarettes and electronic cigarettes^30^. HF is considered an index of parasympathetic modulation of the heart rate, and LF is considered an index of sympathetic modulation, but the actual contribution of the sympathetic and parasympathetic systems to HRV parameters in normal and abnormal physiology are still being dissected^26^. Nevertheless, it is accepted that an increase in the LF to HF ratio of HRV is indicative of sympathovagal disbalance, and usually correlates with development of arrhythmias and poor cardiovascular outcome^26, 31^. This is in line with the in vivo VT inducibility studies which we conducted, where in vaped mice, inducible VT was more sustained compared to air control mice, and where in optical mapping, vaping resulted in action potential instability which manifested as action potential alternans.

In conclusion, our study is the first to show that vaping produces cardiac electrophysiological toxicity that include prolongation of the QT interval, reduced heart rate variability, inducibility of VT, and action potential alternans. The respiratory system is the major route of ENDS smoke entry into the body, and vaping related pulmonary illnesses are increasingly documented in the clinic^37^. However, similarly to combustible tobacco smoking, ENDS use could have potentially serious and harmful effects on the heart. Our work is the first demonstration that vaping compromises the cardiac electrophysiological integrity, and further studies are needed to further assess the long-term cardiac safety profile of ENDS products.

## ACKNOWLEDGEMENTS

This work was supported in part by National Institutes of Health grants R21HL138064, and R01HL129136 to SN, and AHA postdoctoral fellowship to BC.

## REFERENCES

1. Kuehn B. Youth e-Cigarette Use. Jama-Journal of the American Medical Association. 2019;321:138–138.

2. Prochaska JJ. The public health consequences of e-cigarettes: a review by the National Academies of Sciences. A call for more research, a need for regulatory action. Addiction. 2019;114:587–589.

3. Zhu SH, Sun JY, Bonnevie E, Cummins SE, Gamst A, Yin L and Lee M. Four hundred and sixty brands of e-cigarettes and counting: implications for product regulation. Tobacco control. 2014;23 Suppl 3:iii3–9.

4. Zare S, Nemati M and Zheng Y. A systematic review of consumer preference for e-cigarette attributes: Flavor, nicotine strength, and type. PLoS One. 2018;13:e0194145.

5. Day HR, Ambrose BK, Schroeder MJ and Corey CG. Point of Sale Scanner Data for Rapid Surveillance of the E-cigarette Market. Tob Regul Sci. 2017;3:325–332.

6. Leventhal AM, Stone MD, Andrabi N, Barrington-Trimis J, Strong DR, Sussman S and Audrain-McGovern J. Association of e-Cigarette Vaping and Progression to Heavier Patterns of Cigarette Smoking. Jama-Journal of the American Medical Association. 2016;316:1918–1920.

7. Yu V, Rahimy M, Korrapati A, Xuan Y, Zou AE, Krishnan AR, Tsui T, Aguilera JA, Advani S, Crotty Alexander LE, Brumund KT, Wang-Rodriguez J and Ongkeko WM. Electronic cigarettes induce DNA strand breaks and cell death independently of nicotine in cell lines. Oral Oncol. 2016;52:58–65.

8. Hwang JH, Lyes M, Sladewski K, Enany S, McEachern E, Mathew DP, Das S, Moshensky A, Bapat S, Pride DT, Ongkeko WM and Crotty Alexander LE. Electronic cigarette inhalation alters innate immunity and airway cytokines while increasing the virulence of colonizing bacteria. J Mol Med (Berl). 2016;94:667–79.

9. Crotty Alexander LE, Drummond CA, Hepokoski M, Mathew D, Moshensky A, Willeford A, Das S, Singh P, Yong Z, Lee JH, Vega K, Du A, Shin J, Javier C, Tian J, Brown JH and Breen EC. Chronic inhalation of e-cigarette vapor containing nicotine disrupts airway barrier function and induces systemic inflammation and multiorgan fibrosis in mice. Am J Physiol Regul Integr Comp Physiol. 2018;314:R834–R847.

10. Cervellati F, Muresan XM, Sticozzi C, Gambari R, Montagner G, Forman HJ, Torricelli C, Maioli E and Valacchi G. Comparative effects between electronic and cigarette smoke in human keratinocytes and epithelial lung cells. Toxicol In Vitro. 2014;28:999–1005.

11. Scheffler S, Dieken H, Krischenowski O, Forster C, Branscheid D and Aufderheide M. Evaluation of E-cigarette liquid vapor and mainstream cigarette smoke after direct exposure of primary human bronchial epithelial cells. Int J Environ Res Public Health. 2015;12:3915–25.

12. Rowell TR, Reeber SL, Lee SL, Harris RA, Nethery RC, Herring AH, Glish GL and Tarran R. Flavored e-cigarette liquids reduce proliferation and viability in the CALU3 airway epithelial cell line. Am J Physiol Lung Cell Mol Physiol. 2017;313:L52–L66.

13. Sancilio S, Gallorini M, Cataldi A and di Giacomo V. Cytotoxicity and apoptosis induction by e-cigarette fluids in human gingival fibroblasts. Clin Oral Investig. 2016;20:477–83.

14. Palpant NJ, Hofsteen P, Pabon L, Reinecke H and Murry CE. Cardiac development in zebrafish and human embryonic stem cells is inhibited by exposure to tobacco cigarettes and e-cigarettes. PLoS One. 2015;10:e0126259.

15. Farsalinos KE, Romagna G, Allifranchini E, Ripamonti E, Bocchietto E, Todeschi S, Tsiapras D, Kyrzopoulos S and Voudris V. Comparison of the cytotoxic potential of cigarette smoke and electronic cigarette vapour extract on cultured myocardial cells. Int J Environ Res Public Health. 2013;10:5146–62.

16. Khlystov A and Samburova V. Flavoring Compounds Dominate Toxic Aldehyde Production during E-Cigarette Vaping. Environ Sci Technol. 2016;50:13080–13085.

17. Sassano MF, Davis ES, Keating JE, Zorn BT, Kochar TK, Wolfgang MC, Glish GL and Tarran R. Evaluation of e-liquid toxicity using an open-source high-throughput screening assay. PLoS biology. 2018;16:e2003904.

18. Muthumalage T, Prinz M, Ansah KO, Gerloff J, Sundar IK and Rahman I. Inflammatory and Oxidative Responses Induced by Exposure to Commonly Used e-Cigarette Flavoring Chemicals and Flavored e-Liquids without Nicotine. Frontiers in Physiology. 2018;8.

19. Mikheev VB, Buehler SS, Brinkman MC, Granville CA, Lane TE, Ivanov A, Cross KM and Clark PI. The application of commercially available mobile cigarette topography devices for e-cigarette vaping behavior measurements. Nicotine Tob Res. 2018.

20. Cunningham A, Slayford S, Vas C, Gee J, Costigan S and Prasad K. Development, validation and application of a device to measure e-cigarette users’ puffing topography. Scientific Reports. 2016;6.

21. Kosmider L, Jackson A, Leigh N, O’Connor R and Goniewicz ML. Circadian Puffing Behavior and Topography among E-cigarette Users. Tobacco Regulatory Science. 2018;4:41–49.

22. Claycomb WC, Lanson NA, Jr., Stallworth BS, Egeland DB, Delcarpio JB, Bahinski A and Izzo NJ, Jr. HL-1 cells: a cardiac muscle cell line that contracts and retains phenotypic characteristics of the adult cardiomyocyte. Proceedings of the National Academy of Sciences of the United States of America. 1998;95:2979–84.

23. Eddingsaas N, Pagano T, Cummings C, Rahman I, Robinson R and Hensel E. Qualitative Analysis of E-Liquid Emissions as a Function of Flavor Additives Using Two Aerosol Capture Methods. International journal of environmental research and public health. 2018;15.

24. Yepez VA, Kremer LS, Iuso A, Gusic M, Kopajtich R, Konarikova E, Nadel A, Wachutka L, Prokisch H and Gagneur J. OCR-Stats: Robust estimation and statistical testing of mitochondrial respiration activities using Seahorse XF Analyzer. PloS one. 2018;13:e0199938.

25. Rajab M, Jin H, Welzig CM, Albano A, Aronovitz M, Zhang Y, Park HJ, Link MS, Noujaim SF and Galper JB. Increased inducibility of ventricular tachycardia and decreased heart rate variability in a mouse model for type 1 diabetes: effect of pravastatin. American journal of physiology Heart and circulatory physiology. 2013;305:H1807–16.

26. Shaffer F and Ginsberg JP. An Overview of Heart Rate Variability Metrics and Norms. Front Public Health. 2017;5:258.

27. Jin H, Welzig CM, Aronovitz M, Noubary F, Blanton R, Wang B, Rajab M, Albano A, Link MS, Noujaim SF, Park HJ and Galper JB. QRS/T-wave and calcium alternans in a type I diabetic mouse model for spontaneous postmyocardial infarction ventricular tachycardia: A mechanism for the antiarrhythmic effect of statins. Heart rhythm. 2017;14:1406–1416.

28. Noujaim SF, Kaur K, Milstein M, Jones JM, Furspan P, Jiang D, Auerbach DS, Herron T, Meisler MH and Jalife J. A null mutation of the neuronal sodium channel NaV1.6 disrupts action potential propagation and excitation-contraction coupling in the mouse heart. FASEB J. 2012;26:63–72.

29. Zarzoso M, Rysevaite K, Milstein ML, Calvo CJ, Kean AC, Atienza F, Pauza DH, Jalife J and Noujaim SF. Nerves projecting from the intrinsic cardiac ganglia of the pulmonary veins modulate sinoatrial node pacemaker function. Cardiovasc Res. 2013;99:566–75.

30. Moheimani RS, Bhetraratana M, Peters KM, Yang BK, Yin F, Gornbein J, Araujo JA and Middlekauff HR. Sympathomimetic Effects of Acute E-Cigarette Use: Role of Nicotine and Non-Nicotine Constituents. J Am Heart Assoc. 2017;6.

31. Colhoun HM, Francis DP, Rubens MB, Underwood SR and Fuller JH. The association of heart-rate variability with cardiovascular risk factors and coronary artery calcification: a study in type 1 diabetic patients and the general population. Diabetes Care. 2001;24:1108–14.

32. Tse G, Wong ST, Tse V, Lee YT, Lin HY and Yeo JM. Cardiac dynamics: Alternans and arrhythmogenesis. J Arrhythm. 2016;32:411–417.

33. St Helen G and Eaton DL. Public Health Consequences of e-Cigarette Use. JAMA internal medicine. 2018;178:984–986.

34. Aravamudan B, Kiel A, Freeman M, Delmotte P, Thompson M, Vassallo R, Sieck GC, Pabelick CM and Prakash YS. Cigarette smoke-induced mitochondrial fragmentation and dysfunction in human airway smooth muscle. American journal of physiology Lung cellular and molecular physiology. 2014;306:L840–54.

35. luerman G, Obejero-Paz C, Brown A and Bruening-Wright A. Optimizing Rate Correction of Field Potential Duration, a Biomarker for QT Risk Assessment, in Human Ipsc-Cardiomyocytes. Biophysical J. 2014;106:720.

36. Bertino JS, Jr., Owens RC, Jr., Carnes TD and Iannini PB. Gatifloxacin-associated corrected QT interval prolongation, torsades de pointes, and ventricular fibrillation in patients with known risk factors. Clin Infect Dis. 2002;34:861–3.

37. Mukhopadhyay S, Mehrad M, Dammert P, Arrossi AV, Sarda R, Brenner DS, Maldonado F, Choi H and Ghobrial M. Lung Biopsy Findings in Severe Pulmonary Illness Associated With E-Cigarette Use (Vaping). Am J Clin Pathol. 2019.

